# Male Dimorphism and Emergence Strategies: A Mathematical Model of Protandry in *Fabriciana nerippe* Felder, 1862

**DOI:** 10.1101/2024.10.09.617364

**Authors:** Hidaka Kubo, Shinji Nakaoka, Ryo Yamaguchi

## Abstract

In many butterfly species, males emerge earlier than females as part of a strategy to maximize male reproductive success. Although behavioral ecological studies using mathematical models have been conducted to explain this phenomenon, certain emergence patterns remain unexplained. In the butterfly species *Fabriciana nerippe*, some males emerge at the same time as females, in addition to males that emerge earlier than the females. However, it is unclear what emergence patterns occur in populations with male dimorphism, as observed in this species. In this study, we showed the existence of male body size dimorphism in *Fabriciana nerippe* by conducting a comparative analysis of forewing lengths between males and females. In addition, we developed a comprehensive mathematical model to investigate emergence patterns in the presence of dimorphic males. By introducing a trade-off between large size and early emergence, the model considered a scenario where small early-emerging and large late-emerging males could coexist. Numerical analysis demonstrated the emergence patterns of these two male types with a switch in emergence time. Furthermore, the higher the death rate before emergence, the earlier the emergence switch. These findings suggested that the timing of the switch depends on the death rate and is influenced by environmental factors. This work contributes to ecological and theoretical studies on timing dimorphism in life-history strategies across a broader range of species.

**Highlights:**

- We confirmed the existence of male body-size dimorphism corresponding to emergence timing in *Fabriciana nerippe*.
- We developed a comprehensive model of male emergence patterns incorporateing male dimorphism.
- We assumed a trade-off between early emergence and large body size.
- The timing of male emergence switching is influenced by death rate and reproductive advantage of large males.

## 1. Introduction

Life history refers to the sequence of events during an individual’s lifespan. Understanding life histories has long been a central focus in evolutionary ecology. This is because of the accumulation of evidence suggesting that the observed life histories are optimized phenotypes as a result of natural and sexual selection (Stearns, 1976; Stott et al., 2024). Optimal resource allocation and investment schedules for growth and reproduction have been extensively studied, both theoretically and empirically, in reproductive ecology, revealing that different strategies often emerge as optimal solutions for males and females (Shine, 1989).

Protandry, a prime example of a sex-specific reproductive strategy, is where males emerge or mature earlier than females. This strategy maximizes male mating opportunities and minimizes pre-reproductive mortality in females, particularly in species where females mate once and males can mate multiple times (Fagerström & Wiklund, 1982; Wiklund & Fagerström, 1977; Zonneveld & Metz, 1991). The hypothesis that protandry is an adaptive rather than incidental trait due to size differences between the sexes is supported by experimental studies (Wiklund & Solbreck, 1982). This phenomenon is particularly prevalent in butterflies with non-overlapping generations and has been observed in various organisms (Morbey & Ydenberg, 2001).

Although the evolutionary and ecological significance of reproductive timing is well understood, some exceptions exist. One such exception is observed in *Fabriciana nerippe* in Japan, where some males emerge simultaneously with females (late males), in addition to those emerging earlier (early males). Late males are larger than early males in terms of body size. A similar pattern was observed in Dawson’s burrowing bee, *Amegilla dawsoni* (Alcock, 1997). However, the ecological and evolutionary implications of this male dimorphism remain poorly understood and require further research.

The role of male body size in reproductive success may explain the emerging patterns of male dimorphism. Larger males have a reproductive advantage over smaller males, suggesting a trade-off between the benefits of earlier emergence and larger body size. There is a positive correlation between male body size and mating success in butterfly species (Elgar & Pierce, 1988). Additionally, in species where males are territorial, most mating occurs within territories, and males that successfully occupy territories tend to be larger than those that do not (Rosenberg & Enquist, 1991; Wickman, 1985). Furthermore, the reproductive output may depend on both the number and size of ejaculates received by females (Rutowski et al., 1987; Wiklnnd et al., 1993). Since ejaculate size positively correlates with male body size (Boggs, 1981; Svärd & Wiklund, 1986; Wiklund & Kaitala, 1995), larger males may fertilize more eggs, providing an additional reproductive advantage.

Identifying the genetic basis of dimorphism through genome association studies is challenging in *F. nerippe* because it is a non-model organism with an incompletely understood ecology, and its genome has not been sequenced. Consequently, theoretical studies using mathematical models provide a crucial foundation for exploring the evolution of dimorphism in the life history of *F. nerippe*. Several mathematical models have been developed to explain protandry in species where females mate only once or a few times. (Bulmer, 1983; Fagerström & Wiklund, 1982; Wiklund & Fagerström, 1977; Zonneveld, 1992, 1996; Zonneveld & Metz, 1991). These studies primarily focused on the position of emergence time in males and females. However, Iwasa et al., (1983) this approach was expanded by considering both the timing and shape of the emergence patterns. Their model, which was validated using mark–release–recapture data from the checkerspot butterfly *Euphydryas Editha*, revealed that male emergence patterns deviated from a normal distribution, highlighted the importance of considering emergence pattern shapes when studying protandry.

The aim of the current study was to elucidate the phenomenon of male dimorphism in *F. nerippe*. The three main objectives were to 1) confirm the existence of male body size dimorphism by conducting a comparative analysis of forewing lengths in male and female specimens, 2) develop a comprehensive model of male emergence patterns that incorporated this dimorphism, enabling the investigation of its parameter dependencies, and 3) investigate the hypothesis that large, late-emerging males coexist with their smaller, early-emerging counterparts because of a competitive advantage in mating that outweighs the costs of delayed emergence. Through these investigations, we anticipate valuable insights into the evolution and maintenance of alternative male life history strategies, not only in butterflies but also across a broader range of taxa.

## 2. Materials and Methods

### 2.1. Forewing length comparison

#### 2.1.1. Size measurement

The specimens of *Fabriciana nerippe* were used to measure the forewing length. Each individual was photographed alongside a scale bar and label that recorded the date and location of collection. All specimens used in this study were collected between 2013 and 2016 in Higashisonogi, Nagasaki Prefecture, Japan. Forewing length was measured in each photograph using Fiji software (ImageJ) (version 1.54f Schindelin et al., 2012).

#### 2.1.2. Statistical analysis

Males were divided into two groups: early males collected from June 15 to July 1, and late males collected on July 14. Bartlett’s test was conducted among the three groups (early male, late male, and female) to confirm equal variances. The Tukey−Kramer test was conducted to compare the means among the three groups.

### 2.2. Model

We extended a well-developed game-theoretical model originally proposed by Iwasa et al., (1983) to obtain the emergence pattern when there were two types of males in the population. This model enabled us to solve the distribution of eclosion times, which gave all males an equal number of successful mating trials corresponding to the evolutionary equilibrium strategy developed by Maynard Smith & Price, (1973). The variance in male emergence time resulted from an optimization strategy under natural selection. Although the previous model assumed that some males emerged every day in the earlier phase of the mating season and no males emerged later, we assumed that the later phase had late males. The number of emergence events at time *t* (emergence pattern) of early and late males at time *t*are *g*_1_(*t*) and *g*_2_(*t*), respectively. The number of female emergence events at time *t* is denoted as *f*(*t*). *μ*_*a*1_ and *μ*_*a*2_ represent the death rate after emergence, while *μ*_*b*1_ and *μ*_*b*2_ represent the death rate before emergence of early and late males, respectively.

The number of males present at time *t* is the total number of surviving individuals that emerged before time *t*. By multiplying the number of male emergences at time *x* by the survival rate from time *x* to *t* and summing for *x*, the number of males present at time *t*is

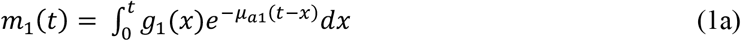

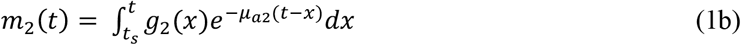

where *t*_*s*_ is the switch time of emergence between the early and late males. We assumed the emergence of male switches at time *t*_*s*_ to simplify the model. Only the early males emerged before *t*_*s*_,, whereas only the late males emerge after *t*_*s*_. The subscripts 1 and 2 refer to early and late males, respectively.

We assumed that females mate immediately after emergence and do not accept males afterward. We also assumed that late males have an advantage over early males in the competition for mate acquisition because of their larger body sizes. Hence, *c* represents the competitive ability of late males over early males (i.e., *c* ≥ 1). The expected number of mates per male at time *t, E*_*i*_(*t*), is:

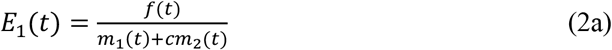

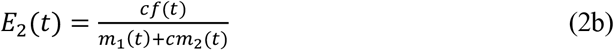

where subscript 1 and 2 refers to early and late males, respectively.

We calculated the fitness of a male emerging on day t: the expected number of matings of a male throughout his life, *ϕ*(*t*). *ϕ*(*t*) is calculated by integrating the expected number of matings per male at time *x, E*(*x*), multiplied by the survival rate from *t* to *x*, and then by the survival rate until emergence.

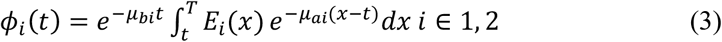

where *T* marks the end of the mating season and *i* denotes the male type (i.e., early or late).

We aimed to determine the form *g*(*t*) takes at equilibrium when the number of that female emergences at time *t, f*(*t*) is fixed, and the number of male emergences at time *t, g*(*t*) evolves,. If males emerging at different times have different fitness levels, the individual with higher fitness will reproduce more offspring, and the form of *g*(*t*) will change. In addition, if the two types of males have different fitness values, the number of males with higher fitness increases, altering the form of *g*(*t*). Therefore, we considered a situation in which the fitness is equal regardless of the emergence time, achieving an evolutionary equilibrium where

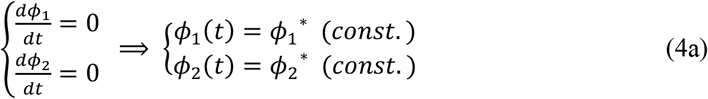

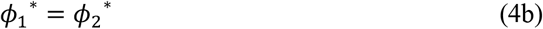

The condition that differentiation of *ϕ*(*t*) = 0 indicated that males that emerged at different times had equal fitness. We assumed that the total number of early and late males that emerged was constant. *M*_*i*_ is the number of type *i* males.

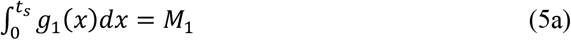

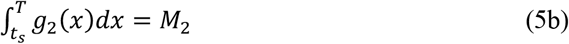

We numerically solved the above equations and explored the parameter dependence of the male emergence curves and the timing of the switch between the two types.

## 3. Results

### 3.1. Forewing length comparison

The results of the Tukey−Kramer test are shown in Fig. 1. This result indicated significant differences in the population means among early males, late males, and females (p < 0.001). Late males had larger forewings than early males, and females had larger forewings than both types of males. The mean forewing lengths of early males, late males, and females were 33.99 ± 0.31, 38.31 ± 0.43, and 41.53 ± 0.72 mm, respectively (mean ± standard error).

**Fig 1.**
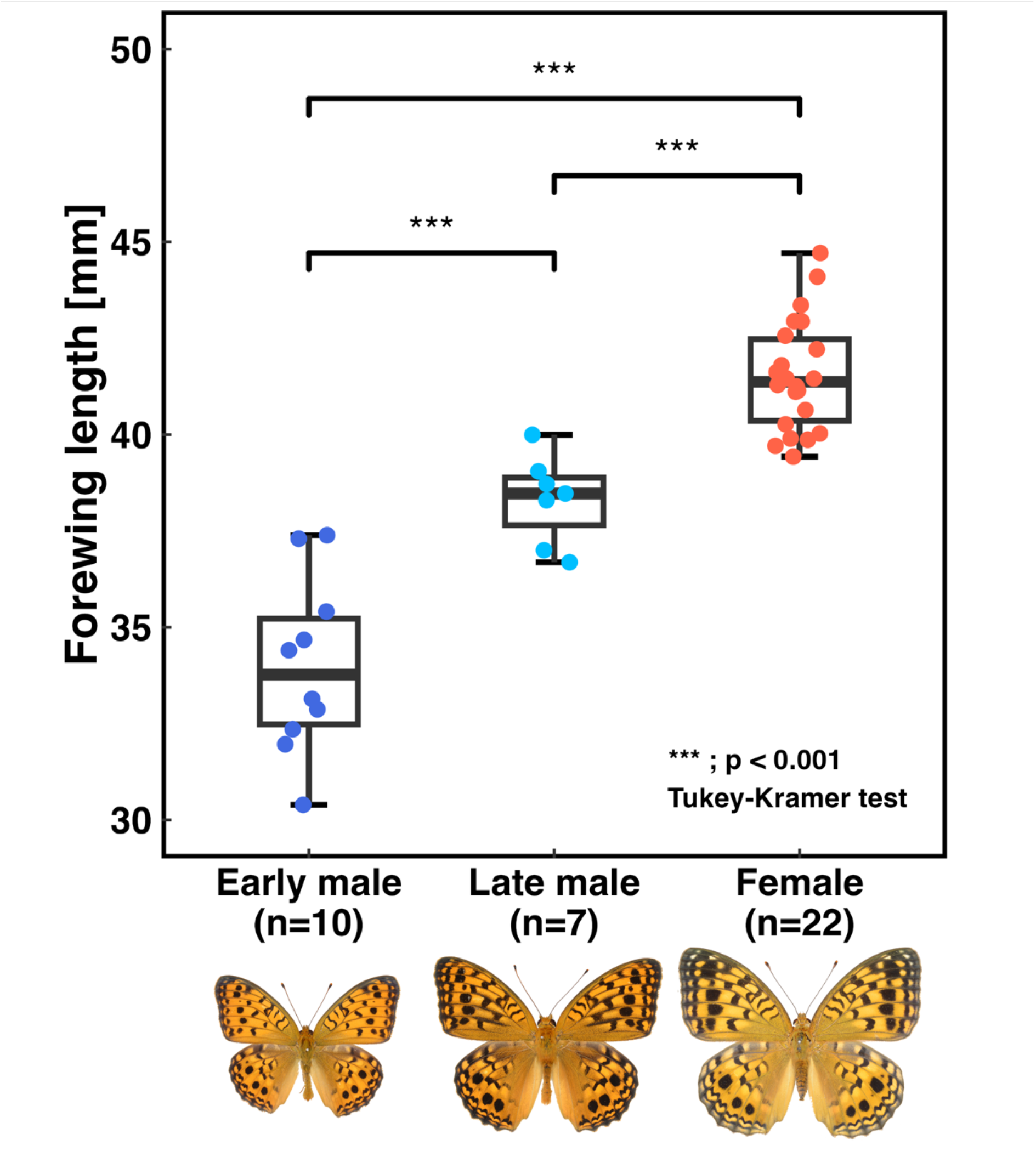
Box plots showing forewing length among early male, late male, and female individuals of *Fabriciana nerippe*. The horizontal axis indicates the categories of data. The vertical axis displays forewing length. The bold horizontal line in each box is the median, and the top and bottom of the box are the first and third quartiles, respectively. The whiskers extend to the maximum and minimum values observed in the data. “n” represents the number of samples in each category. The bars and asterisks at the top of the box represent the result of the Tukey-Kramer test. (*** ; p<0.001). The photographs of the sample individuals shown under each category were taken at the same scale.

### 3.2. Emergence pattern of male butterflies

Fig. 2 shows the emergence pattern of male butterflies when the number of females emerging at time t, *f*(*t*), was followed by the beta distribution. Hereafter, we use *Beta*(5, 3) as the female emergence curve, which is defined in the range [0, 100]. Fig. 2(a) shows that early male emergence transitioned to late male emergence at optimal time *t*_*s*_. Therefore, only early males emerged before *t*_*s*_, and only late males emerged after *t*_*s*_. In this parameter set, *g*_1_(*t*) peaked (local maximum), but *g*_2_(*t*) reached a maximum value at time *t*_*s*_ and monotonically decreased until the end of the mating season. The overall emergence pattern showed that males had an emergence peak earlier than females, indicating the presence of protandry as an evolutionary equilibrium.

**Fig 2.**
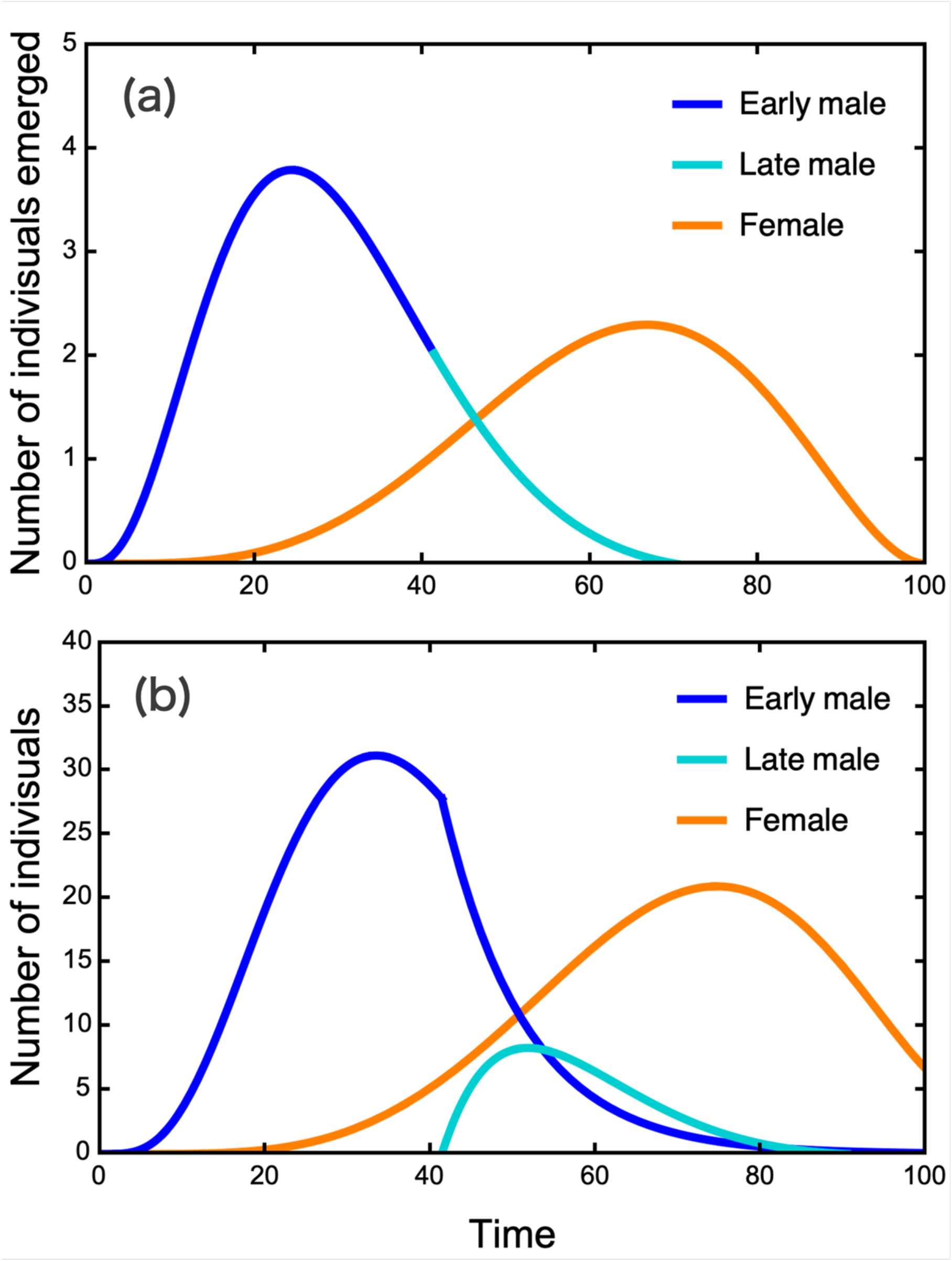
Numerical analysis of butterfly emergence pattern. The horizontal axis indicates time (day). (a) The top graph shows the number of emergences at time *t* and (b) the bottom graph shows the number of individuals at time *t*. Blue, light blue, and orange lines represent early male, late male, and female, respectively. Parameters are *μ*_a1_=0.1, *μ*_a2_=0.1, *μ*_b1_=0.09, *μ* _b2_=0.09, *T*=100, *c*=1.5, *M*_1_=100, and *M*_2_=20.

Fig. 2(b) shows the number of individuals at time t, *m*_1_(*t*), *m*_2_(*t*), and *F*(*t*)(i.e., 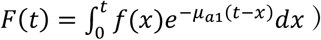. The number of early males decreased exponentially from *t*_*s*_, whereas late males began to emerge from *t*_*s*_, corresponding to the switch in emergence at time *t*_*s*_. The peaks in the number of individuals appeared in the order of early males, late males, and females, indicating that protandry occurred in both males. In this parameter set, there was a period when the late male population was larger than the early male population.

### 3.3. Parameter dependence of *t*_*s*_

We examined the parameter dependence of *t*_*s*_ to investigate features of emergence pattern. Fig. 3 shows the dependence of the switch time *t*_*s*_ on the death rate after emergence *μ*_*a*_ and before emergence *μ*_*b*_. We simply assumed that death rate after emergence *μ*_*a*_ is larger than death rate before emergence *μ*_*b*_. Thus, simulations in this figure, were performed in the range of 0.01 ≤ *μ*_*b*_ ≤ 0.09 and 0.1 ≤ *μ*_*a*_ ≤ 0.95. The larger the death rate after emergence *μ*_*a*_, the later *t*_*s*_. In contrast, the larger the death rate before emergence *μ*_*b*_, the earlier *t*_*s*_. *t*_*s*_ increased linearly with *μ*_*b*_, while the rate of increase of *t*_*s*_ gradually decreased with *μ*_*a*_. In addition, *μ*_*b*_ has a larger effect on *t*_*s*_ than *μ*_*a*_, even though the range of *μ*_*b*_ was smaller than that of *μ*_*a*_. Therefore, *μ*_*b*_ had a greater effect on *t*_*s*_ than *μ*_*a*_.

**Fig 3.**
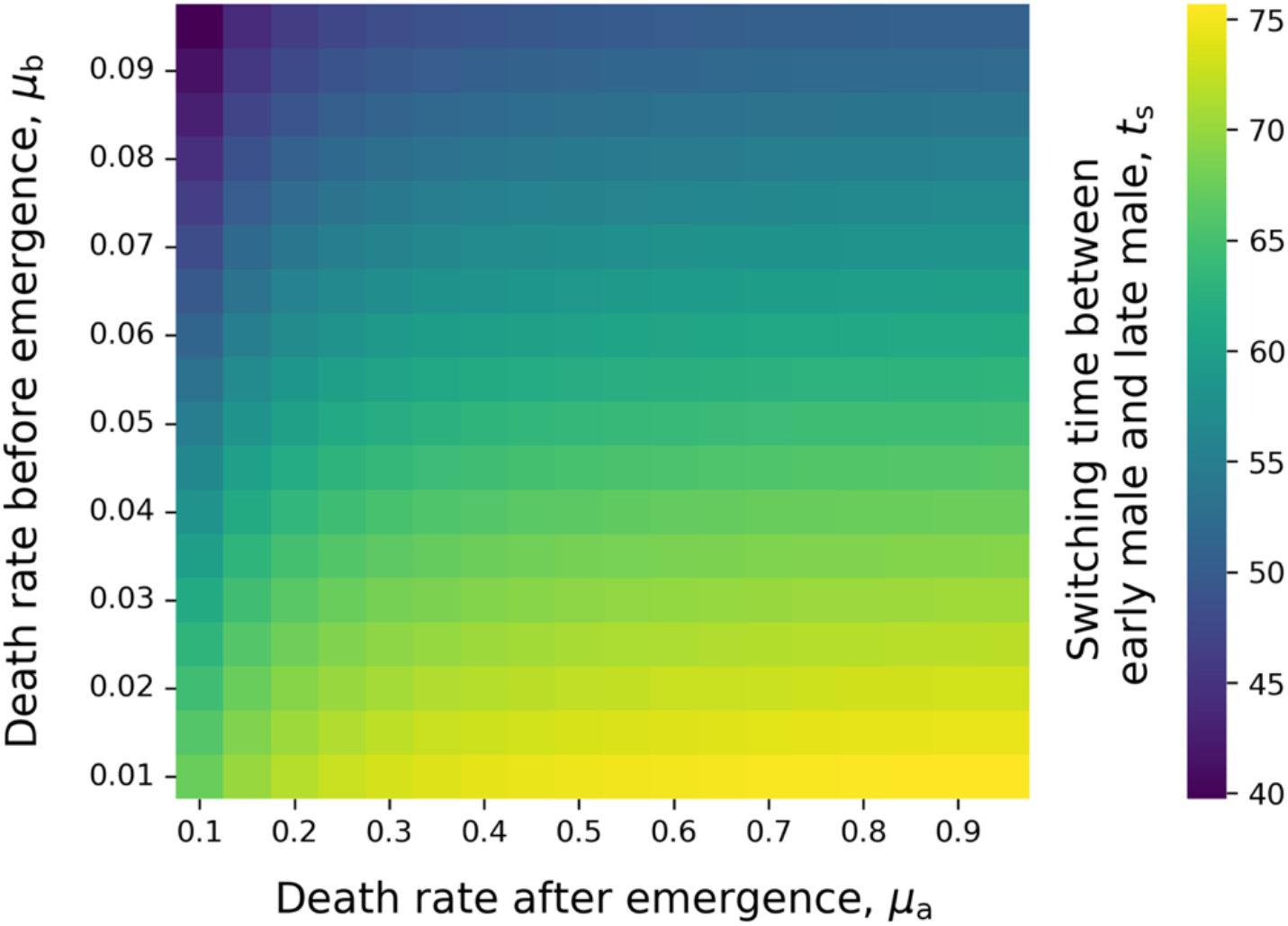
Dependence of switching time *t*_s_ on the death rate after emergence *μ*_a_ and before emergence *μ*_b_. Dark regions indicate early switching time, while light-colored regions indicate later switching time. Other parameters are *T*=100, *c*=1.5, *M*_1_=100, *M*_2_=20.

Fig. 4 shows the dependence of the switch time *t*_*s*_ on the competitive advantage of late male *c* . The results of simulations with three different *M*_2_ values are shown. As *t*_*s*_ decreased with *c* for any *M*_2_, the late male advantage increased, and the switch time occurred earlier. When *M*_2_ assumed a large value, *t*_*s*_ was small. In addition, the change in *t*_*s*_ owing to *M*_2_ was larger than that owing to *c*.

**Fig 4.**
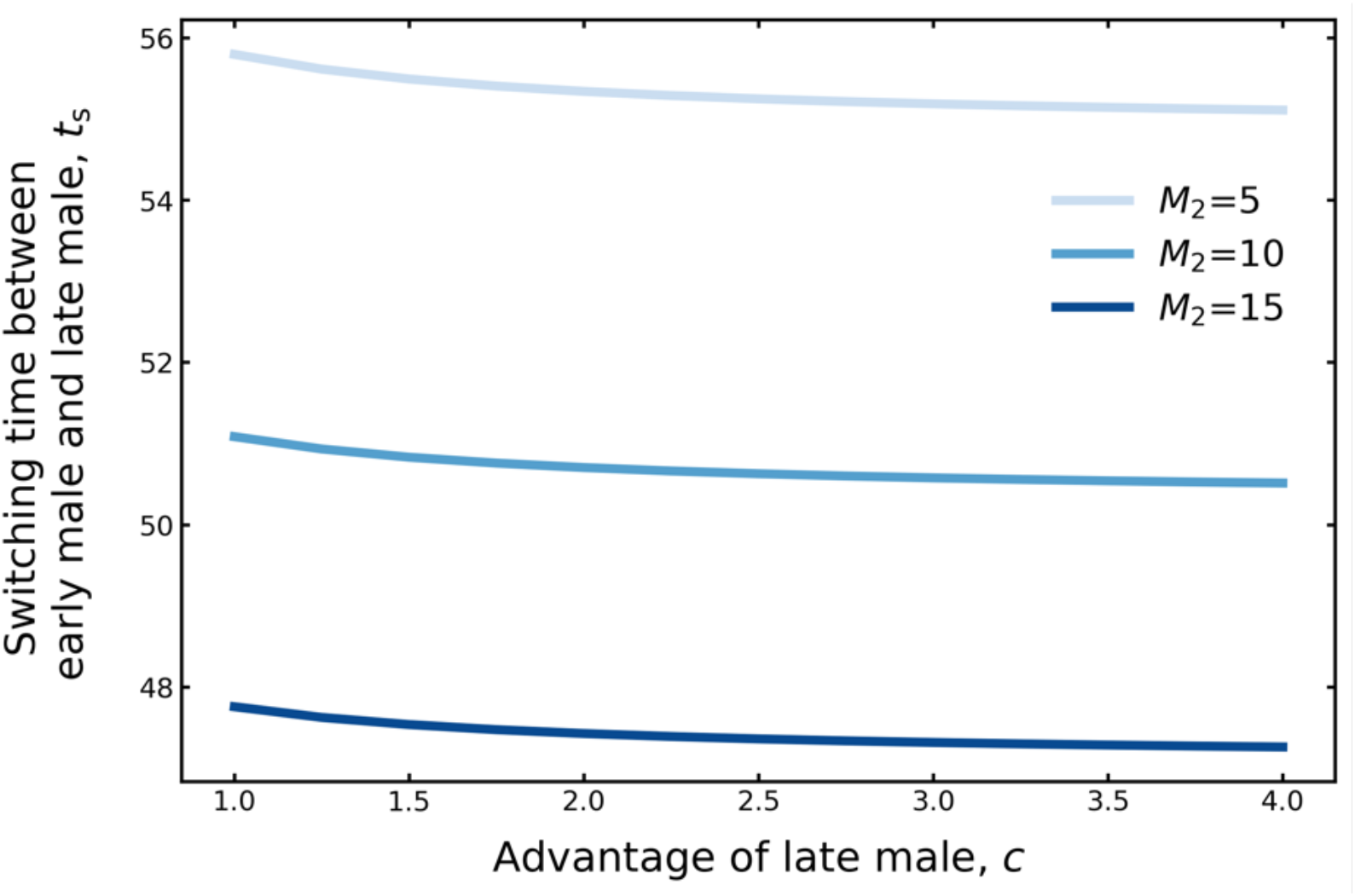
Dependence of switching time *t*_s_ on the competitive advantage of late male *c*. Cases with three different total numbers of late male emergence (*M*_2_=5, 10, and 15) are shown. Total number of all male emergence *M*_1_+*M*_2_ is fixed at 100. Other parameters are *μ*_a1_=0.11, *μ*_a2_=0.1, *μ*_b1_=0.09, *μ*_b2_=0.09, *T*=100.

## 4. Discussion

This study presents a significant advancement in our understanding of protandry and male dimorphism in butterflies, specifically in *F. nerippe*. We empirically confirmed the known male body size dimorphism in this species (Fig. 1) and extended the classic mathematical model of protandry to account for the emergence patterns of males when male dimorphism exists in a focal population (Fig. 2). Although previous studies on protandry have not considered dimorphism in the population (e.g., Iwasa et al., 1983; Wiklund & Fagerström, 1977), we focused on the shape of the male emergence pattern when the male was dimorphic in body size and emergence timing, representing a novel approach in behavioral ecology.

Our model revealed that mortality rates influenced emergence timing. Higher pre-emergence mortality led to earlier switch times, whereas higher post-emergence mortality delayed switch (Fig. 3). As spending a longer period in a life stage with higher mortality leads to lower fitness, the species may have evolved to spend less time in this stage. In addition, the effect of the death rate before emergence on ts is greater than that of the death rate after emergence. This suggested that environmental factors that affect mortality rates, such as climate change or invasive species, can significantly affect emergence patterns. We also found that a greater advantage for late-emerging males *c* consistently resulted in earlier switch times (Fig. 4). This trend did not change for any of the combinations of parameters explored. Larger, late-emergence males had a greater advantage when competing with early males than when competing with late males. This insight provides a new perspective on the evolution of emergent strategies in dimorphic populations.

Various studies in the fields of reproductive and behavioral ecology have focused on dimorphism in reproductive traits within a sex (Gross, 1996). Polymorphism in the reproductive behavior of ruff (Hogan-Warburg, 1965; Jukema & Piersma, 2006) and marine isopods (Shuster, 1989) is a well-known example. In addition to the above examples, many other alternative reproductive traits have been identified, most of which focus on body features and behavior. Our work sheds light on the previously overlooked aspects of temporal dimorphism in life history strategies, particularly emergence timing, in *F. nerippe*. Our description of male dimorphism in *F. nerippe* and the accompanying mathematical modeling will pave the way for future ecological and theoretical studies on timing dimorphism in life history strategies across broader species.

Field studies collecting time-series emergence data and detailed mortality rates across life stages are crucial to further validate our model. A previous protandry model in butterflies was validated using time-series emergence count data obtained using the mark-and-recapture method and wing damage observations (Baughman et al., 1988; Iwasa et al., 1983; Sawada et al., 1961). Validation of the model requires not only daily abundance and size measurements but also time-series abundance data or emergence count data. Acquiring such detailed field data could reveal additional polymorphisms in emergence timing, similar to the ones we focused on in *F. nerippe*, and significantly deepen our understanding of these traits across species.

Moreover, precise mortality data are crucial for refining this model. We incorporated both adult and pre-emergence mortalities (during the larval and pupal stages) into our calculations. This approach aligned with the established knowledge that death rates vary significantly across different butterfly life stages (Baker, 1970; Moore, 1989). Future field studies and model development should carefully consider the stage-specific mortality rates to enhance the accuracy and predictive power of protandry models.

Several simplifying assumptions made in this study warrant further investigation. First, a key assumption of our model is the abrupt switch between early and late male emergence at a specific time. However, empirical studies have shown that emergence patterns typically lack clear truncation at onset or conclusion (Baughman et al., 1988; Iwasa et al., 1983; Sawada et al., 1961). This discrepancy may be explained by stochastic environmental factors, microhabitat variations (Iwasa et al., 1983), or interannual fluctuations in male mortality rates (Iwasa & Haccou, 1994). Future models could be enhanced by incorporating these factors to better reflect the gradual nature of the emergence patterns observed in natural populations.

Furthermore, our model implicitly assumed a genetic basis for male dimorphism, with both male types having equal fitness. While this genetic determination of dimorphism has been observed in species such as ruffs, marine isopods, and Batesian mimetic butterflies (Iijima et al., 2018; Küpper et al., 2015; Shuster & Sassaman, 1997), other organisms such as earwigs and dung beetles exhibit status-dependent dimorphisms in behavior and phenotype (Emlen, 1994; Tomkins & Brown, 2004). Future modeling efforts should consider the potential effects of environmentally dependent male dimorphism on population dynamics. For instance, a recent study on round gobies (Young et al., 2023) demonstrated that environmentally determined male tactics, specifically the presence of sneaker males, can increase overall population size.

A critical question that remains is the evolutionary origin of dimorphism in *F. nerippe*. Our study modeled emergence patterns under the assumption of existing dimorphism; however, the underlying evolutionary mechanisms require further investigation. Adaptive dynamics (Geritz et al., 1998; Metz et al., 1992) provides an effective framework for studying the conditions under which dimorphism evolves from monomorphism. This approach allows a theoretical description of evolutionary branching within the context of an evolutionarily stable strategy. Furthermore, it enables the exploration of not only the emergence of dimorphism but also the evolution of the proportion of dimorphic individuals within a population. Among the fritillaries (subgenus *Fabriciana* of the genus *Argynnis*), only *F. nerippe* exhibits dimorphism in the timing of male emergence. This raises important questions for future research. Why is this dimorphism absent in other closely related species? Which specific conditions promote the evolution of dimorphism in terms of emergence timing and body size? Addressing these questions will deepen our understanding of *F. nerippe* and contribute to broader theories of life history, evolution, and reproductive strategies in insects.

## Acknowledgement

We are grateful to Dr. Y. Tachiki for his insightful discussions. We also extend our sincere thanks to the staff of the Kitakyushu Museum of Natural History & Human History for providing access to the butterfly specimens. We would like to thank to Nakaoka lab for their invaluable discussions and support throughout this research. This work was supported by funding from the JSPS KAKENHI (24K02092 to RY).

## Code availability

The codes that produce the figures in this study is written in Mathematica and Python, and these will be available on Github (https://github.com/hidakakubo/e_pattern) upon acceptance.

